# Moderate Cold Stress Enhance Drought Resistance through *CCA1* via -an ABA-independent Pathway

**DOI:** 10.1101/2024.07.09.602734

**Authors:** Xue Yang, Yan Liu, Zi-Chang Jia, Ming Li, Xuan-Xuan Hou, Sheng-Qiang Hou, Xi-Long Shi, Bei Gao, Dao-Yuan Zhang, Fu-Yuan Zhu, Mo-Xian Chen, Ying-Gao Liu

**Affiliations:** State Key Laboratory of Green Pesticide; Key Laboratory of Green Pesticide and Agricultural Bioengineering, Ministry of Education, Center for R&D of Fine Chemicals of Guizhou University, Guiyang, 550025, China; State Key Laboratory of Crop Biology, College of Life Science, Shandong Agricultural University, Tai’an 271018, China; Co-Innovation Center for Sustainable Forestry in Southern China, College of Biology and the Environment, Nanjing Forestry University, Nanjing, China; State Key Laboratory of Desert and Oasis Ecology, Xinjiang Institute of Ecology and Geography, Chinese Academy of Sciences, Urumqi 830011, China

**Keywords:** Cold stress, Drought tolerance, *CCA1*, *OST1*, *P5CS1*, ABA independent

## Abstract

In nature, plants frequently encounter concurrent stresses, particularly the simultaneous occurrence of cold and drought stress poses a challenge to plants in middle and high latitudes. However, the molecular mechanisms underlying the plants response to this double-stress scenario remain unclear. Although some responses suggest that drought stress can improve cold resistance in plants, through ABA signaling pathways. In our study, we discovered that moderate low temperature treatment significantly enhanced Arabidopsis drought tolerance. Low temperature rapidly triggers the transcription factor CCA1, a prototypical response to cold stress, which inturn directly regulates the expression of *OST1* and *P5CS1* by binding to their promoters. This leads to the premature closure of stomata and accumulation of proline through a non-ABA-dependent pathway even before plants experience drought stress, ultimately improving plant resistance against drought and cold. Moreover, this mechanism is conserved across plant species, and the synergistic resistance mechanism enables perennial plants to survive winter conditions and annual plants to withstand multi-stresses.

## Introduction

Low temperature and drought are two abiotic stresses that affect plant growth and development and limit the quality and yield of many crops in the world (Chen et al. 2021). At present, research on plant low temperature and drought stress alone is relatively mature(Gupta, Rico-Medina, and Caño-Delgado 2020; Shi, Ding, and Yang 2018b). But in nature, most of the time drought and cold stress will occur at the same time, especially in winter of Temperate continental climate and temperate monsoon climate (Lu et al. 2020), and it is an important factor limiting agricultural productivity and affecting perennials over winter (Jiang et al. 2017; Wang et al. 2016). The multiple genomics analysis and genome-wide association research showed that there were overlaps of key transcripts, proteins and metabolites in plants under low temperature stress and drought stress (Xu and Fu 2022; Shinozaki and Yamaguchi-Shinozaki 2022). The observation also implies the possibility of a significant signal overlap between cold.

There is a notable crossover in how plants respond to low temperature and drought stress, underlining the intricate interconnection of these environmental challenges(Olson et al. 2018). Low temperature and drought stress can lead to osmotic stress. Among them, the osmotic stress signal can be sensed by calcium ion channel OSCA1(reduced hyperosmolality-induced calcium increase 1) (Yuan et al. 2014) and mechanical sensitive channel Msc S-like protein MSL8 (Hamilton et al. 2015). Osmotic stress results in the increase of calcium ions, which is related to the cyclic nucleotide-gated ion channel (CNGCs) family and glutamate-like receptor (GLR) family (Swarbreck, Colaço, and Davies 2013). Both cold and drought stress affect membrane proteins on cells, including various transporters and ion channels, as well as membrane receptor kinases (RLKs) (Sangwan et al. 2002; Liu et al. 2017). Both kinds of stress can promote the production of ROS such as superoxide anion, hydrogen peroxide, hydroxyl radical and singlet oxygen, seriously affect the ROS control system, and produce a series of secondary signal molecules (Mignolet-Spruyt et al. 2016). The two stresses also accumulate stress hormone ABA, and the accumulated ROS and ABA act as signaling molecules to regulate stress response (He et al. 2012; Wang et al. 2018). Among them, the calcineurin B-like protein-CBL interacting protein kinase (CPL-CIPK) signaling pathway in plants is a Ca²+ correlated pathway that responds strongly to both low temperature and drought stress (Yu, An, and Li 2014). Under low temperature and drought stress, MAPK signaling pathway studies observed multiple MAPKs, mainly MPK3, 4 and 6, which activated downstream target genes in response to stress (de Zelicourt, Colcombet, and Hirt 2016; Li et al. 2017). Calcium-regulated protein kinases such as CPKs and CBL-CIPKs and Raf kinases activate the SnRK2 family of protein kinases (Fàbregas, Yoshida, and Fernie 2020; Lin et al. 2020), which then activate abscisic acid, the osmoregulatory substance proline, and the second messenger IP3 (Saruhashi et al. 2015; Fujii, Verslues, and Zhu 2011; Ding et al. 2018a; Ding et al. 2018b; Takahashi et al. 2020). Both stressors trigger overlapping signaling pathways and gene expression patterns in plants, often involving stress-related transcription factors, hormones and metabolic changes (sugar and amino acid accumulation) (Eckardt 2019; Zhao et al. 2022; Li et al. 2023; Gong et al. 2020; Zhou and Zhang 2020). Additionally, various protective molecules, such as like proline and antioxidants, are commonly induced in response to both stresses, showcasing the shared adaptive strategies plants employ to mitigate damage caused by drought and low temperature (Guo et al. 2021; Jin et al. 2023). Understanding this shared response is pivotal for devising comprehensive strategies to enhance plant resilience to these stressors and ensure sustainable crop yields.

Spring sowing corn is often affected by low temperature, drought and other harsh environment. Transcriptomic studies on maize germination under cold and drought stress showed that TFs, hormone metabolism and signal transduction, flavonoid metabolism, sucrose metabolism and cell growth promotion were involved in drought resistance (Li et al. 2021). According to the phylogenetic analysis and expression patterns of TCP transcription factors in cassava seedlings under cold and drought stress, the expression patterns of MeTCPs change under cold and drought stress, and ABA signaling pathway plays a key role in this response (Lei et al. 2017). According to the genome-wide identification and expression results of PP2C gene of M. runcatula under drought and cold stress, *MtPP2C46*, *MtPP2C47* and *MtPP2C72* can be up-regulated by cold and drought induction, while the expression of *MtPP2C92* under drought and cold treatment is reversed (Yang et al. 2018). Studies showed that cold and low temperature stress activated the expression of GIBBERELLIN SUPPRESSING FACTOR (GSF), and GSF inhibited the synthesis of GA by inhibiting *GA20oxs* and activating the expression of *GA2oxs*, thereby slowing down plant growth and improving the stress resistance (Chen, Li, and Yang 2019). Overexpression of *OsSCL30* resulted in increased accumulation of reactive oxygen species and decreased resistance of transgenic rice to low temperature, drought and salt stress(Zhang, Sun, et al. 2022). Through genome-wide identification and expression analysis of LEA protein gene family in tea plant, LEA gene is involved in low temperature and drought stress (Jin et al. 2019).

Current scientific research results show that pretreatment of plants with drought stress can effectively improve tolerance to various abiotic stresses, including low temperature. In the nutritional stage of drought stress, spring wheat enhances its tolerance to low temperature stress by improving its antioxidant capacity and photosynthetic efficiency (Li et al. 2015; Mahajan and Tuteja 2005). Even the freeze resistance of perennial ryegrass (sensitive variety) is improved by drought pretreatment and increased carbohydrate, proline and protein contents (Hoffman et al. 2012). In chickpea, preexposure to nonlethal drought stress can improve plant resistance to low temperature by reducing membrane damage and oxidative stress (Saini et al. 2019). ABA treatment before plant exposure to cold stress improves maize survival under cold stress, which proves that endogenous ABA activated by drought pretreatment plays an important role in plant adaptation to cold (Guo et al. 2021).

However, it is still unknown whether cold stress has an impact on plant drought resistance and how cold stress affects plant drought resistance. Our study demonstrates that moderate low temperature treatment significantly enhances the drought resistance of plants. Additionally, low temperature treatment rapidly triggers the expression of CIRCADIAN CLOCK ASSOCIATED 1 (*CCA1*), subsequently elevating the expression levels of OPEN STOMATA 1 (*OST1*) and DELTA1-PYRROLINE-5-CARBOXYLATE SYNTHASE 1 (*P5CS1*). This process enables early closure of stomata and accumulation of proline, a crucial resistant substance, even before plants experience drought stress. Such synergistic mechanisms play a critical role in enabling plants to cope with multiple stresses, particularly for overwintering species in aridity and low temperatures regions. This research provides guidance for the breeding and genetic modification of crops, aiming to enhance their resilience against multiple challenging stresses, ultimately leading to improved agricultural productivity and food security.

## Results

### Moderate cold treatment promoted the adaptation of Arabidopsis thaliana to drought stress

In order to investigate the impact of moderate cold treatment (12℃) on drought tolerance in Arabidopsis thaliana, 3-week-old seedlings were subjected to a 14-day treatment involving moderate cold, drought, and both drought and moderate cold. Compared to the control group, moderate cold had a slight effect on growth speed after the 14-day treatment period, while drought significantly impacted growth speed (Figure 1A). Under dual stress conditions of drought and moderate cold, although plant growth was weaker compared to only moderate cold treatment, it was significantly better than that under only drought treatment (Figure 1A). This suggests that moderate cold stress aids plants in resisting drought stress. Additionally, several physiological characteristics of Arabidopsis plants were measured across different treatment groups. The results revealed that cold treatment had a minor influence on relative water content, fresh weight, survival ratio, MDA levels, electrical conductivity and proline contents of plants; however, drought treatment had a significant impact on these parameters. Furthermore, dual stress greatly mitigated the effects of solely undergoing drought treatment on relative water content, survival ratio, MDA levels, proline contents, fresh weight and relative electrical conductivity (Figure 1, B-E, Supplemental Figure S1H-1I). These findings indicate moderate cold treatments enhance Arabidopsis adaptability to withstand drought stress.

**Figure 1.**
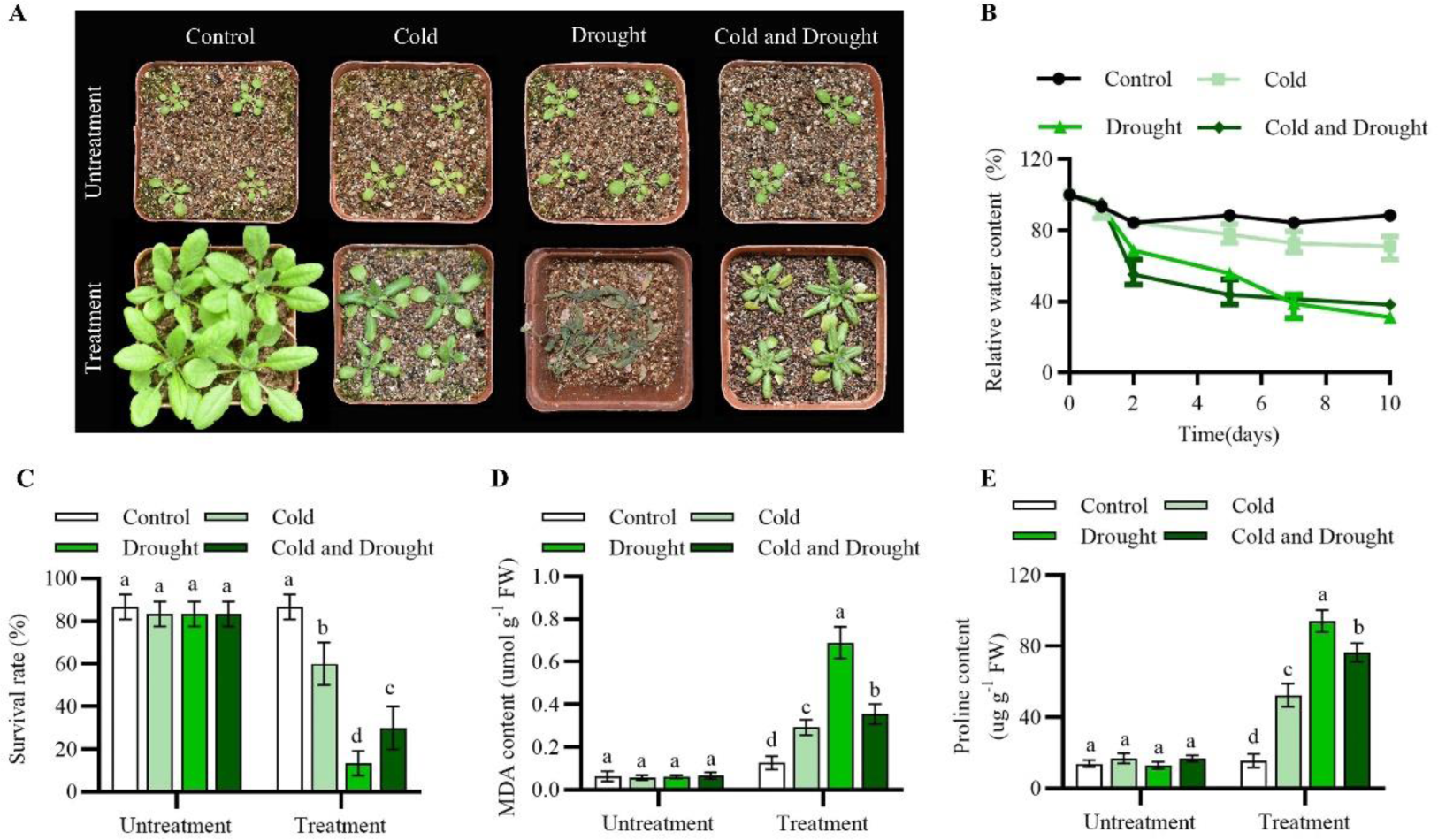
Moderate cold stress functions in drought tolerance in *Arabidopsis*. A, Phenotypic analysis of *Arabidopsis* seedlings under cold and drought stress. Three-week-old Arabidopsis plants were exposed to 12°C and a soil moisture content of 80% for 14 days (control), to 12°C and a soil moisture content of 80% for 14 days (cold), to 23°C and a soil moisture content of 40% for 14 days (drought), and to 12°C and a soil moisture content of 40% (cold and drought). B, Statistical analysis of the relative water content of the seedlings in (A). C, Statistical analysis of the survival rate of the seedlings in (A). D, Statistical analysis of the malondialdehyde (MDA) content of the seedlings in (A). E, Statistical analysis of the proline content of the seedlings in (A). The data are presented as the means ± SEMs (n = 3 biological replicates). Different letters indicate significant differences (*P* < 0.05, one-way ANOVA, Tukey’s test).

We also investigated the impact of intense low temperature stress (4℃) on Arabidopsis, and our findings demonstrate that both growth and development of plants are significantly influenced by exposure to 4℃ cold treatment. Moreover, the combined stress phenotype resulting from simultaneous exposure to 4℃ cold treatment and drought treatment is more severe compared to individual stresses of either drought or cold alone (Supplemental Figure S1A-1G). Collectively, these results suggest that moderate rather than strong cold stress contributes to enhancing plant resilience against drought stress.

### The response to cold stress was earlier than that to drought stress

To investigate the mechanism by which moderate cold stress enhances drought resistance in *Arabidopsis thaliana*, we analyzed the expression changes of moderate cold-induced and drought-induced genes under different treatment conditions. Our findings demonstrate that both moderate cold stress and dual treatment of cold and drought stress can rapidly induce the expression of INDUCER OF CBF EXPRESSION 1 (*ICE1*), *CCA1*, and other genes. Specifically, *ICE1* and *CCA1* was significantly and rapidly (0.5 h) induced as a response to cold stress, and prolonged exposure to moderate cold stress could induce the expression of drought-responsive genes such as RESPONSIVE TO ABA 18 (*RAB18*) and RESPONSIVE TO DESICCATION 26 (*RD26*) (Figure 2). However, drought stress alone did not induce relative genes expression at early adversity, it took at least two days to induce the expression of these drought-stress-related genes (ABA DEFICIENT 1 (*ABA1*), ABA INSENSITIVE 1 (*ABI1*), *RAB18* and *RD26*), and there was no significant effect on the expression of genes related to cold stress until the end (Figure 2, E-H). The findings suggest that exposure to moderate cold stress can rapidly and significantly trigger the expression of genes associated with both cold stress and drought, with the response to drought stress notably lagging behind that of cold stress.

**Figure 2.**
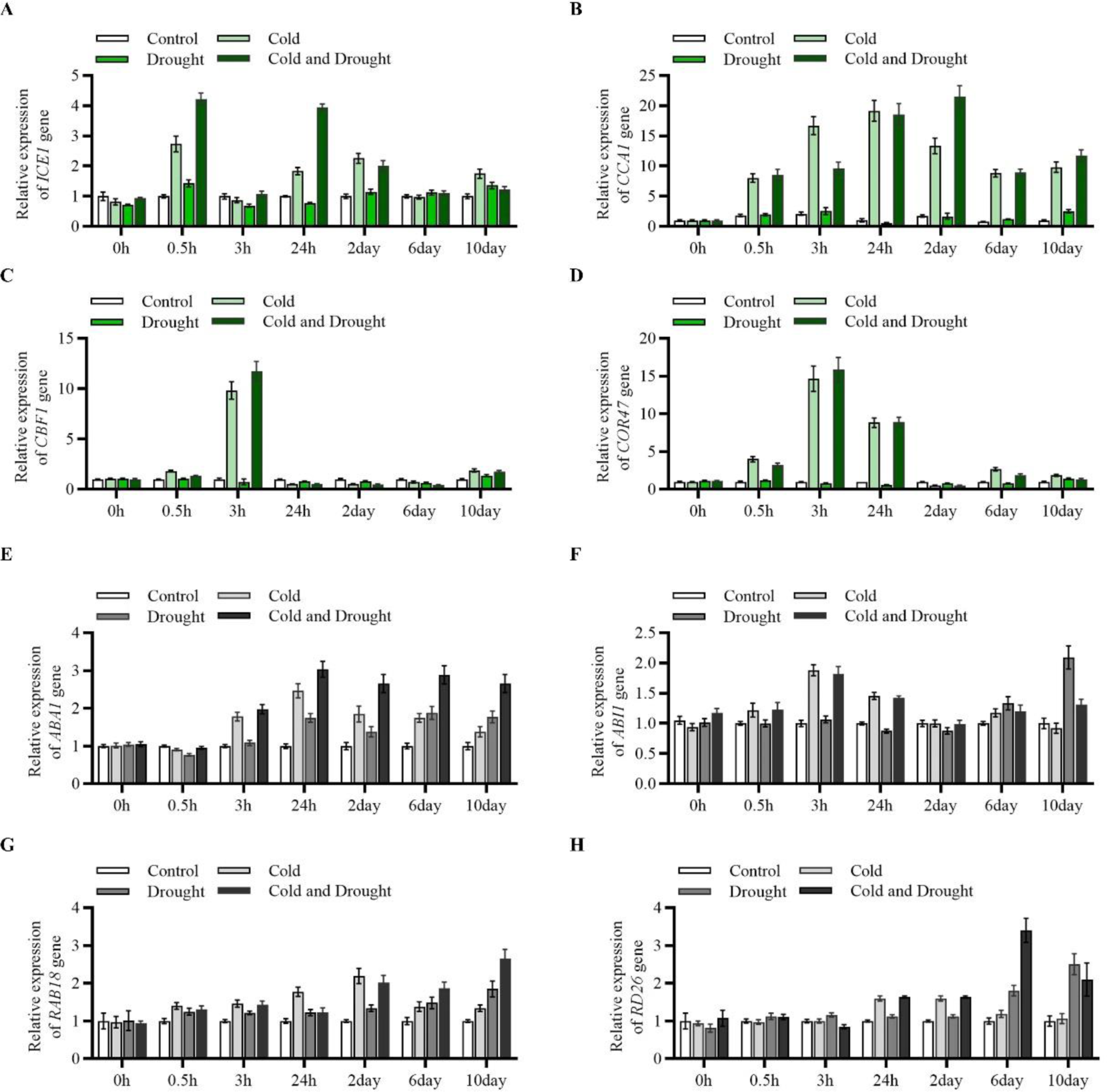
The response to cold stress occurred earlier than that to drought stress in *Arabidopsis thaliana*. Relative expression levels of *ICE1* (A), *CCA1* (B), *CBF1* (C), *COR47* (D), *ABA1* (E), *ABI1* (F), *RAB18* (G) and *RD26* (H) in Col0 plants under CK, cold, drought, cold and drought stress conditions.

Since the response to moderate cold stress occurred significantly earlier than that to drought stress, and moderate cold stress exhibited a beneficial effect on drought stress, we postulated that certain cold-induced transcription factors might regulate the expression of drought stress-related genes. Therefore, we investigated the expression patterns of transcription factors associated with key responses to cold stress under various treatment conditions. The results demonstrated significant induction of all these transcription factors by cold stress, and albeit to a lesser extent by drought stress simultaneously (Supplemental Figure S2A). Notably, *CCA1* and *ICE1* were exclusively induced by moderate cold stress but not by drought stress, particularly *CCA1* which was only induced by moderate cold (Supplemental Figure S2B). Consequently, *CCA1* was selected as a potential candidate gene for subsequent investigations.

### *CCA1* is required for cold-induced drought resistance

The core gene of plant circadian rhythm, *CCA1* (Hsu and Harmer 2014), plays multiple roles in addition to its regulation of circadian rhythm, including involvement in autophagy, fatty acid synthesis, and the response to cold stress in plant cells (Chen et al. 2022; Kim et al. 2023; Kidokoro et al. 2021). To investigate the function of *CCA1* under cold and drought stress conditions, we obtained *CCA1* knock-down and over-expressed plants (Supplemental Figure S2, E-H).

As a transcription factor, CCA1 is localized in the nucleus (Supplemental Figure S2C). *CCA1* was expressed in all tissues, especially in cauline leaves (Supplemental Figure S2D). Compared to the wild type, *cca1* mutants exhibited significantly heightened susceptibility to drought and moderate cold stress, whereas *CCA1* over-expressing plants demonstrated markedly enhanced resistance (Figure 3A). After 14 days dual stress treatment, the experiment on relative water content demonstrated a significantly lower water content in *cca1* compared to the wild type, whereas the over-expression of *CCA1* exhibited a significantly higher water content than the wild type (Figure 3B). The results of fresh content after treatment also confirmed the above results (Figure 3C). We also conducted tests on the stomatal aperture of various materials, and the results confirmed that the stomatal aperture of *cca1* was significantly higher than that of the wild type under dual stress. Conversely, over-expression of *CCA1* led to a significant reduction in stomatal aperture compared to the wild type (Figure 3, D and E). The results also showed that *CCA1-OE* strains accumulated more proline after dual stress treatment (Figure 3F). And *cca1* mutants exhibited increased accumulation of MDA and higher relative conductivity, whereas *CCA1* over-expressing strains displayed lower relative conductivity and reduced MDA levels (Figure 3, G and H). These findings indicate that drought and moderate cold dual stress induce significant membrane damage in *cca1* mutants, while *CCA1* overexpressing strains experience significantly less membrane damage.

**Figure 3.**
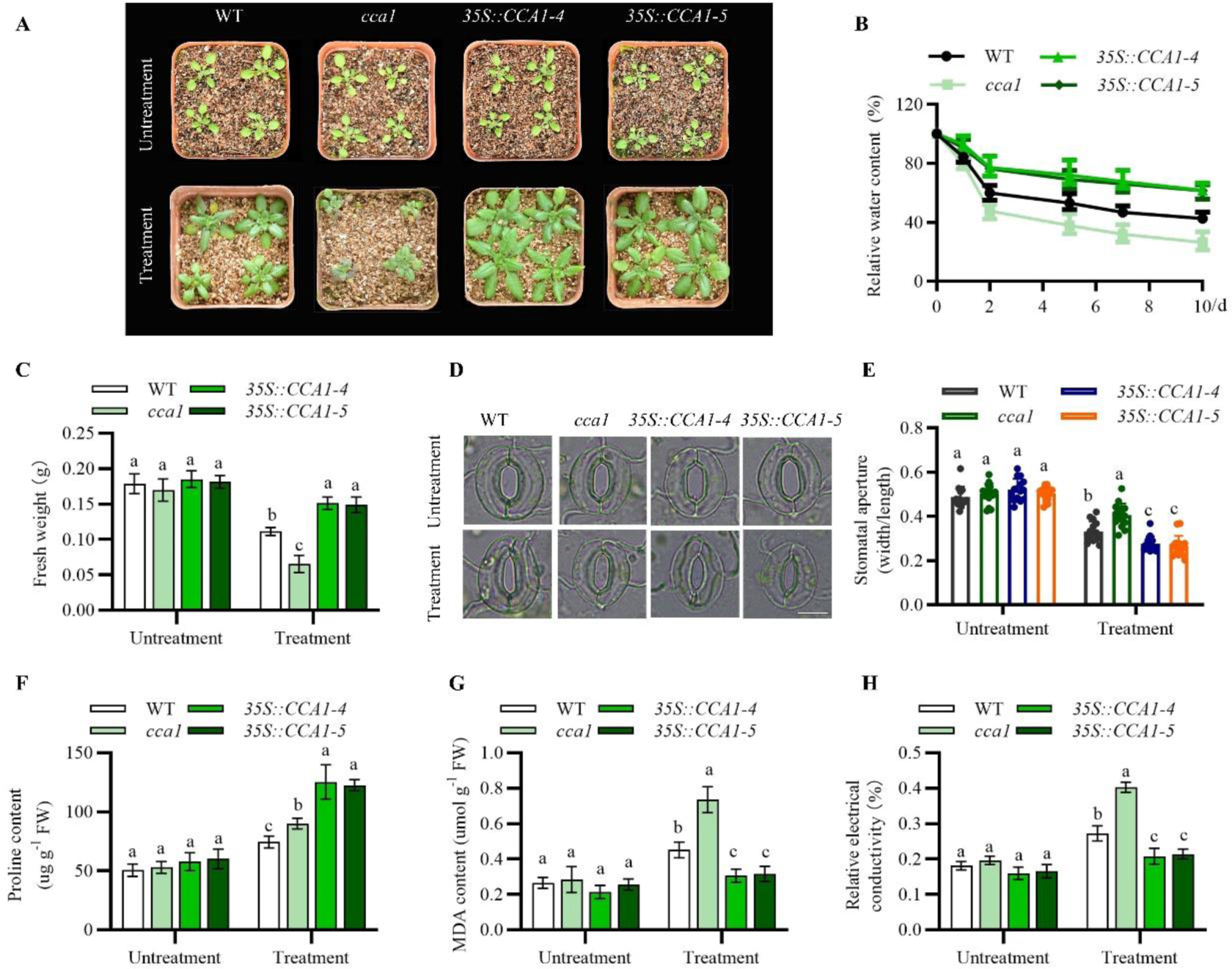
*CCA1* functions in drought tolerance under cold stress in *Arabidopsis*. A, Representative seedlings of the WT, *cca1, 35S::CCA1*-4, and *35S::CCA1*-5 plants subjected to 12°C and 40% soil moisture content treatment on the 14th day. *At*WT is the Columbia ecotype. B, Statistical analysis of the RWC of the samples in (A). C, Statistical analysis of the fresh weight of the samples in (A). D, Stomatal phenotype analysis of the WT, *cca1, 35S::CCA1*-4, and *35S::CCA1*-5 plants under drought and cold stress. Photographs were taken on the 7th day after seed sowing. Scale bars, 20 µm. E, Statistical analysis of the stomatal aperture of the samples in (D). F, Statistical analysis of the relative electrical conductivity of the samples in (A). G, Statistical analysis of the MDA content of the samples in (A). H, Statistical analysis of the proline content of the samples in (A). The data are presented as the means ± SEMs (n = 3 biological replicates). Different letters indicate significant differences (*P* < 0.05, one-way ANOVA, Tukey’s test).

The changes in gene expression in response to drought and cold stress were also monitored. The results demonstrated a significant induction of genes associated with the response to cold stress, such as C-REPEAT/DRE BINDING FACTOR 1 (*CBF1*), *CBF2*, and *CBF3*, during the early stage of treatment (3h), which was further enhanced in lines overexpressing *CCA1* (Supplemental Figure S3, A-C). However, there was no significant induction observed for drought-related genes like *ABA1*, *ABI1* and *RAB18* (Supplemental Figure S3, D-F). Interestingly, we observed that the expression of drought-related *P5CS1* (associated with proline synthesis) and *OST1* (associated with stomatal closure) genes was induced by moderate cold stress and double stress in early treatment. However, these genes were not induced by drought stress alone. Moreover, this induction was significantly enhanced in *CCA1* overexpressing lines but completely abolished in cca1 mutants (Supplemental Figure S3, G and H). These findings suggest that cold stress may activate the expression of certain drought response genes through *CCA1*.

### The CCA1-regulated cold stress-facilitated drought stress response requires the involvement of *P5CS1* and *OST1*

To further explore the roles of CCA1 and identify the output genes, we collected the ChIP-seq data of CCA1 from multiple studies (Adams et al. 2018; Nagel et al. 2015; Kamioka et al. 2016). We then overlapped these target genes with the Predicted CCAl-binding genes and drought genes. This led to the identification of 10 genes (Supplemental Figure S4, A), that are supposed to be output genes of CCA1. Notably, *P5CS1* and *OST1* were among the 10 genes, consistent with our findings that CCA1 may be involved in multiple stress processes and suggest the potential role of *P5CS1* and *OST1* in regulating multiple stress (Supplemental Figure S4, B). To further understand how CCA1 binds to DNA sequences of *P5CS1* and *OST1*, we used AlphaFold3 (Abramson et al. 2024) to generate structural models of protein-DNA interactions. Utilizing AlphaFold3’s algorithms, we have generated a three-dimensional model of CCAl-pro*P5CS1* and CCAl-pro*OST1* (Supplemental Figure S4, C and D). The results suggest that CCA1, as a transcription factor, can directly bind to the promoter regions of *P5CS1* and *OST1*.

To further validate the ability of the transcription factor CCA1 to specifically bind to the promoter *P5CS1* and *OST1* and regulate their expression, Dual-Luciferase Reporter assay in *N. benthamiana* leaves verified that CCA1 activates the expression of *P5CS1* and *OST1*. In vivo LUC fluorescence showed that CCA1 activated the expression of *P5CS1* and *OST1* (Figure 4, A and D). The results of relative LUC activity measured by double luciferase also support this result (Figure 4, B and E). Then, we first used JASPAR to predict multiple *P5CS1* and *OST1* promoter motifs that CCA1 may bind to, and finally, we selected “AAAATATCT” as the probe and labeled biotin for electrophoretic mobility shift assay (EMSA) experiments. The resutls of EMSA showed that probe-pro*P5CS1*-CCA1 binding complexes and probe-pro*OST1*-CCA1 binding complexes were detected (Figure 4, C and F). We also analyzed the promoter of *P5CS1* and *OST1* and identified a major binding site of CCA1, Evening Element (EE) segment (AAATATC) (Figure 4, G and I). The EE is a major cis-acting element in the promoters of CCA1 and is known to be a CCA1 binding site (Kidokoro et al. 2021; Kamioka et al. 2016; Nagel et al. 2015). ChIP (Chromatin immunoprecipitation) assays confirmed that CCA1 can bind the promoter of *P5CS1* and *OST1* in vivo (Figure 4, H and J). Then, we used microscale thermophoresis (MST) to quantify the binding affinity of CCA1 to the DNA sequences of *P5CS1* and *OST1*, respectively. The resultant data sets demonstrated distinct binding affinities between CCA1 and the promoters of *P5CS1* and *OST1* (Figure 4, K and L), indicating that there is a significantly strong binding affinity between the two components. These results suggest that CCA1 can bind to the promoter of *P5CS1* <u>and</u> *OST1* and regulate their expression.

**Figure 4.**
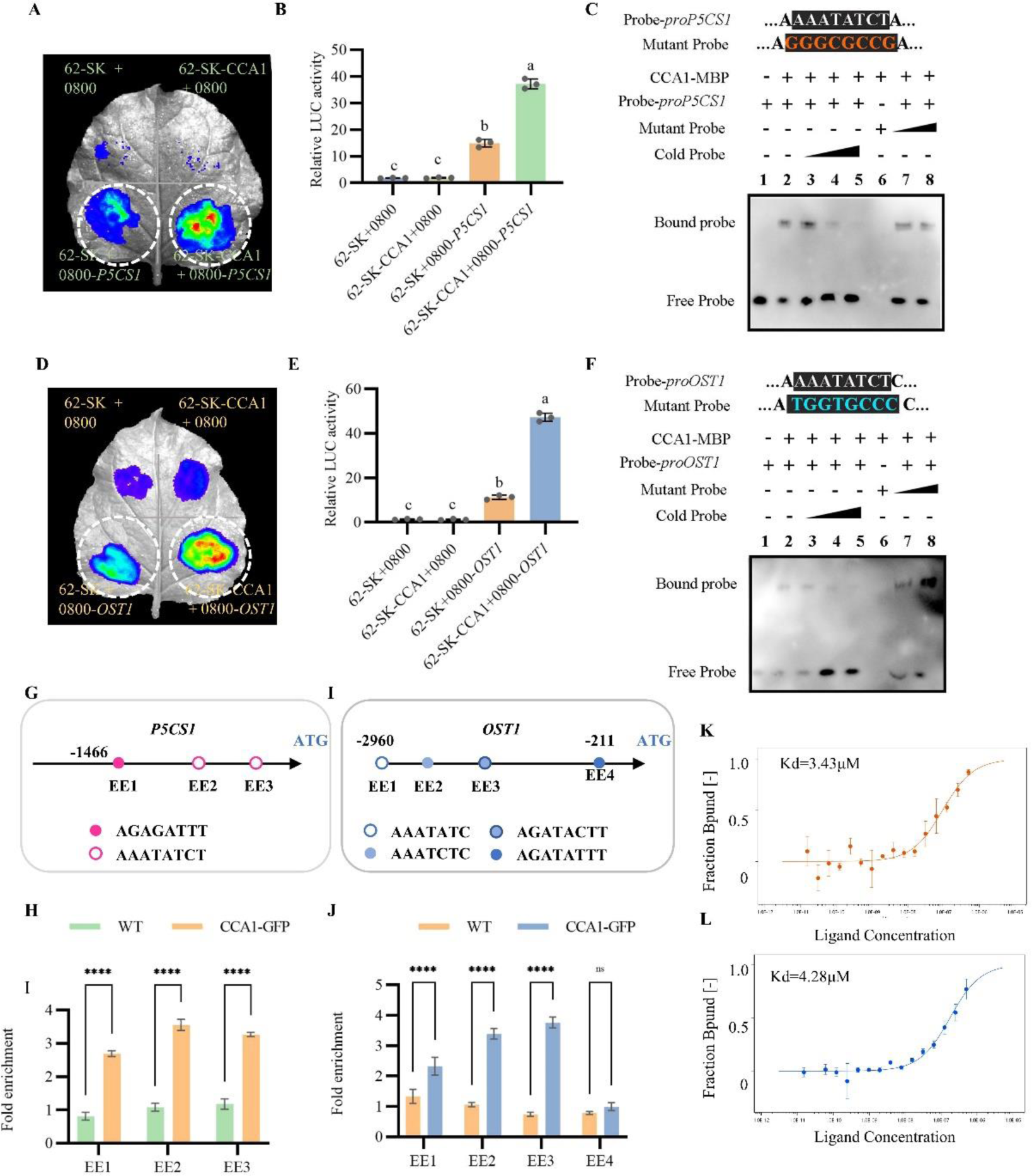
CCA1 promotes the expression of *OST1* and *P5CS1*, which function in plant drought tolerance under cold stress. A, Luciferase reporter experiment of both CCA1 and the promoter of *P5CS1*. B, Statistical analysis of the relative fluorescence activity in (A). C, EMSA of both CCA1 and the hot probe of *P5CS1*. Cold and mutant probes were used as competitive probes. D, Luciferase reporter experiment of both CCA1 and the promoter of *OST1*. E, Statistical analysis of the relative fluorescence activity in (D). F, EMSA of both CCA1 and the hot probe of *OST1*. Cold and mutant probes were used as competitive probes. G Motif analysis of the *P5CS1* promoter. The 1466 bp upstream region of *P5CS1* was analysed. The red ball indicates AGAGATTT, a segment of EE. Red circles indicate AAATATC, a segment of EE. EE1 to EE3 indicate primer pairs used for ChIP‒qPCR in (H). H, CCA1 binds to the promoter of *P5CS1* in vivo. Twelve-day-old Col-0 and CCA1-GFP plants were harvested for ChIP assays. Chromatin fragments were immunoprecipitated with anti-GFP beads (IP) or without immunoprecipitation (input). The precipitated DNA was analysed by RT‒qPCR using the primer pairs indicated in (G). The level of binding was calculated as the ratio between the IP and the input, normalized to that of *Actin8* as an internal control. Error bars, SDs of three biological replicates. *P < 0.05, by t test using Excel. I, Motif analysis of the *OST1* promoter. The 2960 bp region upstream of *OST1* was analysed. The blue balls indicate AGATACTT, AATCTC, and AGATATTT, which are segments of the EE. The blue circles indicate AAATATC, a segment of EE. EE1 to EE4 indicate primer pairs used for ChIP‒qPCR in (J). J, CCA1 binds to the promoter of *OST1* in vivo. Twelve-day-old Col-0 and CCA1-GFP plants were harvested for ChIP assays. Chromatin fragments were immunoprecipitated with anti-GFP beads (IP) or without immunoprecipitation (input). The precipitated DNA was analysed by RT‒qPCR using the primer pairs indicated in (I). The level of binding was calculated as the ratio between the IP and the input, normalized to that of *Actin8* as an internal control. Error bars, SDs of three biological replicates. *P < 0.05, by t test using Excel. K-L, MST analysis of CCA1 binding directly to the *P5CS1* promoter (K) and *OST1* promoter (L). The data are presented as the means ± SEMs (n = 3 biological replicates). Different letters indicate significant differences (*P* < 0.05, one-way ANOVA, Tukey’s test).

Next, we performed genetic experiments to verify that CCA1 may activate *P5CS1* and *OST1* expression under cold stress, thus allowing plants to withstand drought stress. we identified the T-DNA insertion mutants SALK_063517C (*p5cs1*) (Supplemental Figure S5, A and B) and SALK_008068C (*ost1*) (Supplemental Figure S5, E and F). We overexpressed *CCA1* in *p5cs1* and *ost1* plants and identified *35S::CCA1/p5cs1* mutant (Supplemental Figure 5, C and D) and *35S::CCA1/ost1* mutant (Supplemental Figure 5, G and H), respectively. On untreament condition, the phenotype of WT, *35S::CCA1*, *35S::CCA1/p5cs1* and *p5cs1* mutant were consistent. Moreover, after dual stress, overexpression of *CCA1* in *p5cs1* mutants not rescued stress tolerance caused by *p5cs1* deletion (Figure 5A). Moreover, Overexpression *CCA1* mutants further lines exhibited enhanced resistance. *P5CS1* mediates drought resistance in plants by regulating proline accumulation (Shrestha et al. 2022). The proline content determination experiment showed that the *35S::CCA1/p5cs1* accumulated proline at lower levels similar to the *p5cs1* mutant and the *35S::CCA1* line displayed elevated proline levels comparable to WT (Figure 5B). These results suggest that *P5CS1* may act downstream of CCA1 on drought tolerance. The results of fresh weight, MDA content and relative ion content all confirmed this conclusion (Figure 5, C-E). In addition, the phenotype of WT, *35S::CCA1*, *35S::CCA1/ost1* and *ost1* mutant were consistent under normol condition. Moreover, after dual stress, overexpression of *CCA1* in *ost1* mutants not rescued stress tolerance caused by *ost1* deletion (Figure 5F). Moreover, Overexpression *CCA1* mutants further lines exhibited enhanced resistance. *OST1* mediates drought resistance in plants by regulating leads to stomata closure (Upadhyay 2022). The stomatal aperture assay showed that the *35S::CCA1/ost1* maintained a high level of stomatal opening similar to the *p5cs1* mutant and the stomatal aperture of the *35S::CCA1* line displayed significantly reduced under stress comparable to WT (Figure 5, H and I). These results suggest that *OST1* may act downstream of CCA1 on drought tolerance. The results of MDA content and relative ion content all confirmed this conclusion (Figure 5, G and J). Taken together, these results suggest that CCA1 regulates drought tolerance by affecting the *P5CS1* and *OST1* expression under cold stress.

**Figure 5.**
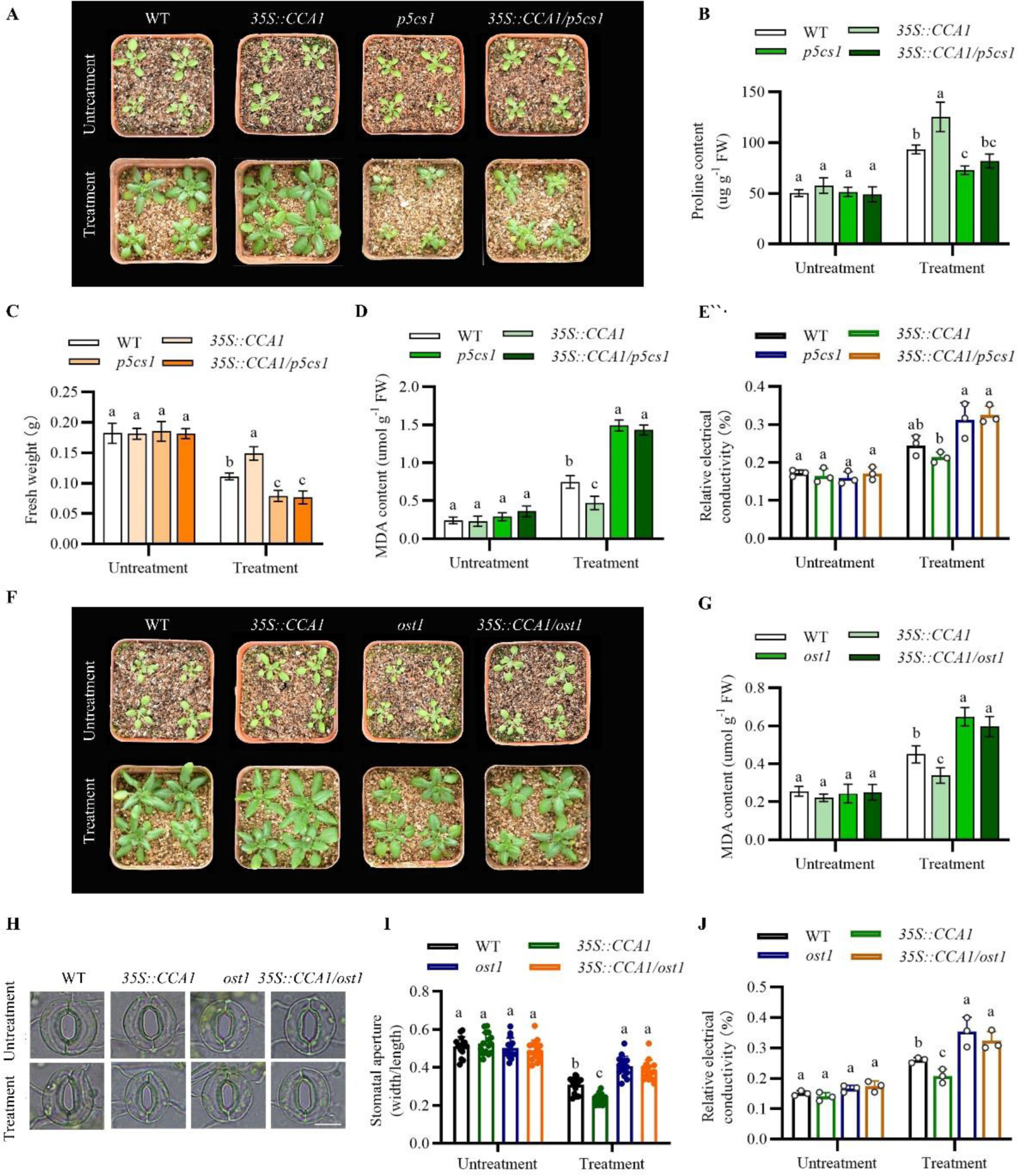
*CCA1* functions in plant drought tolerance under cold stress by promoting the expression of *OST1* and *P5CS1*. A, Representative seedlings of the WT, *35S::CCA1*, *p5cs1* and *35S::CCA1/p5cs1* plants subjected to 12°C and 40% soil moisture content (cold and drought) treatments on the 14th day. B, Statistical analysis of the proline content of the samples in (A). C, Statistical analysis of the fresh weight of the samples in (A). D, Statistical analysis of the MDA content of the samples in (A). E, Statistical analysis of the relative electrical conductivity of the samples in (A). F, Representative seedlings of the WT, *35S::CCA1*, *ost1* and *35S::CCA1/ost1* plants subjected to 12°C and 40% soil moisture content (cold and drought) treatment on the 14th day. G, Statistical analysis of the MDA content of the samples in (F). H, Stomatal phenotype analysis of *At*WT, *AtCCA1-OE*, *Atost1* and *AtCCA1/ost1* under drought and cold stress. Scale bars, 20 µm. I, Statistical analysis of the stomatal aperture of the samples in (H). J, Statistical analysis of the relative electrical conductivity of the samples in (F). The data are presented as the means ± SEMs (n = 3 biological replicates). Different letters indicate significant differences (*P* < 0.05, one-way ANOVA, Tukey’s HSD test).

### CCA1 enhances cold-induced drought resistance in an ABA-independent manner

As a stress hormone, ABA is involved in the regulation of various abiotic resistances, and it has been reported that cold stress can induce an increase in ABA content (de Zelicourt, Colcombet, and Hirt 2016; Waadt et al. 2022; Ding et al. 2015). Therefore, we wanted to further investigate whether the regulation of *P5CS1* and *OST1* by CCA1 requires the involvement of ABA. Then we monitored the association between ABA accumulation and stomatal opening under different treatment conditions using fluorescent staining techniques. The results showed that stomatal closure was earlier than ABA accumulation under cold treatment (Figure S6).

Then the *aba1* mutant was subjected to screening (Supplemental Figure S7, A and B), and subsequently, the *35S::CCA1/aba1* double mutant was constructed (Supplemental Figure S7, C and D). The phenotypic results demonstrated that over-expression of *CCA1* enhanced plant resistance to dual stress, and *aba1* mutants showed sensitive to dual stress. Notably, the over-expression of *CCA1* in the *aba1* mutant significantly compensated for the diminished resistance phenotype (Figure 6A). The results of MDA content indicated that the over-expression of *CCA1* in the *aba1* background significantly alleviated the stress level of *aba1* (Figure 6B). The stomatal aperture assay showed that the *35S::CCA1/aba1* maintained a lower level of stomatal opening similar to the *35S::CCA1* lines and the stomatal aperture of the *aba1* mutant line displayed significantly high stomatal opening under stress comparable to WT (Figure 6, C and D). The proline content determination experiment showed that the *35S::CCA1/aba1* accumulated proline at high levels similar to the *35S::CCA1* line and the *aba1* mutant displayed reduced proline levels comparable to WT (Figure 6E). We also investigated the ABA content in various materials under different treatment conditions using ABA activity staining. The findings revealed that, after 3h treatment dual stress elicited a slight induction of ABA accumulation in wide type (Supplemental Figure S7E). Results also showed that *35S::CCA1* lines did not accumulate more ABA. However, ABA remained at a lower concentration in the *aba1* and *35S::CCA1/aba1* double mutant (Supplemental Figure S7E). The findings suggest that *CCA1* has the ability to regulate the expression of *P5CS1* and *OST1* via an ABA-independent pathway.

**Figure 6.**
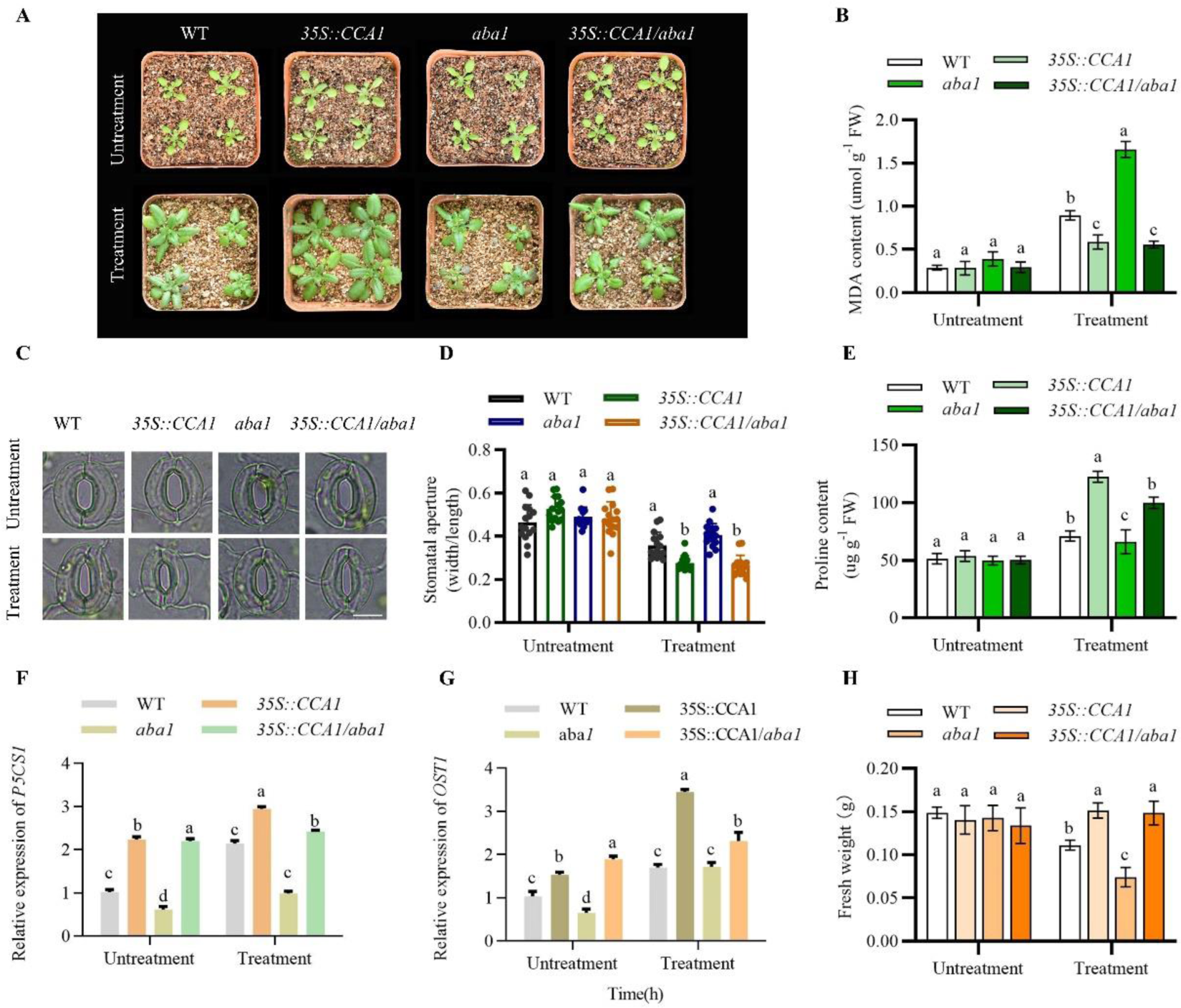
*CCA1* regulates the plant drought response under cold stress through ABA-independent pathways. A, Representative seedlings of the WT, *35S::CCA1*, *aba1* and *35S::CCA1/aba1* plants subjected to 12°C and 40% soil moisture content (cold and drought) treatment on the 14th day. B, Statistical analysis of the MDA content of the samples in (A). C, Stomatal phenotype analysis of WT, *35S::CCA1*, *aba1* and *35S::CCA1/aba1* plants under drought and cold stress. Scale bars, 20 µm. D, Statistical analysis of the stomatal aperture of the samples in (C). E, Statistical analysis of the proline content of the samples in (A). F, Relative expression levels of *P5CS1* in the WT, *35S::CCA1*, *aba1* and *35S::CCA1/aba1* plants after treatment. G, Relative expression levels of *OST1* in the WT, *35S::CCA1*, *aba1* and *35S::CCA1/aba1* plants after treatment. H, Statistical analysis of the fresh weight of the samples in (A). The data are presented as the means ± SEMs (n = 3 biological replicates). Different letters indicate significant differences (*P* < 0.05, one-way ANOVA, Tukey’s HSD test).

The expression of the corresponding ABA-responsive genes, except for *P5CS1* and *OST1*, was not significantly increased in *35S::CCA1*, and all of these genes were significantly decreased in *aba1*, and dule treatment also could slightly increased the expression of *P5CS1* and *OST1* in *aba1* (Figure 6, F and G; and Supplemental Figure S7, F-I). However, with the exception of *P5CS1* and *OST1*, all ABA-responsive genes exhibited significant down-regulation in *35S::CCA1/aba1* double mutants, whereas *P5CS1* and *OST1* displayed up-regulation (Figure 6, F and G). The findings suggest that *CCA1* has the ability to regulate the expression of *P5CS1* and *OST1* via an ABA-independent pathway, thereby influencing cold-induced drought resistance.

### Moderate low temperature pretreatment improved plant cold resistance

To investigate the effect of moderate cold treatment (12°C) on cold tolerance in *Arabidopsis thaliana*, 3-week-old plants were subjected to moderate cold treatment, direct cold treatment (4°C), or moderate cold pretreatment for 2 days prior to cold treatment (Supplemental Figure S8E). The results revealed that moderate cold had a minimal impact on the growth rate after 14 days of treatment compared to the control group. However, a significant reduction in damage caused by sharp cold treatment was observed with a 2-day moderate cold pretreatment (Figure 7A), as evidenced by higher survial rate and lower MDA content in pretreated plants (Figure 7, B and D). The results also showed that moderate low temperature treatment and pretreatment could significantly improve the expression level of CCA1 (Figure 7C).

**Figure 7.**
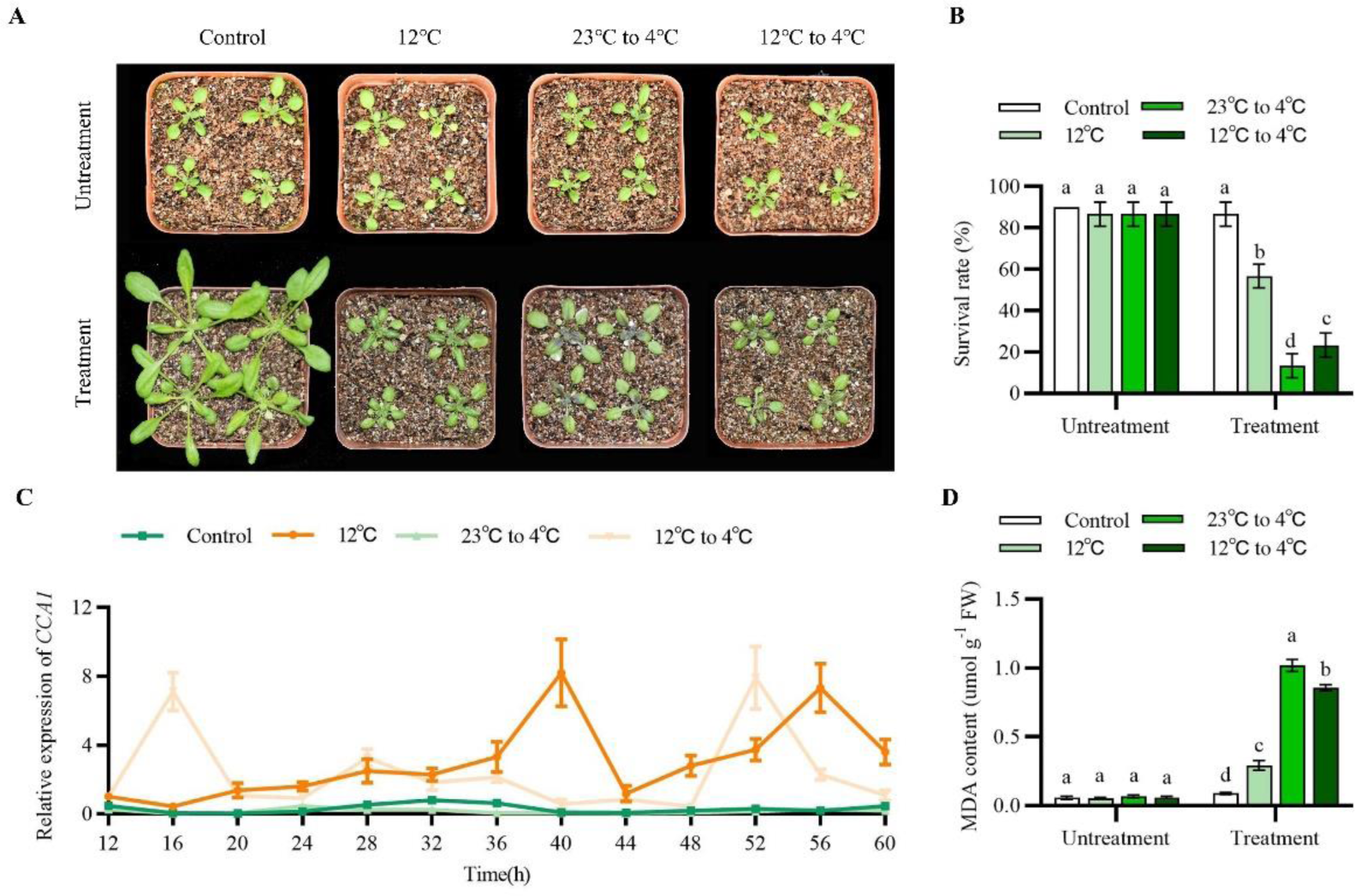
Moderate low-temperature pretreatment improved plant cold resistance. A, Phenotypic analysis of seedlings under different cold stresses. Three-week-old *Arabidopsis plants* were exposed to 23°C, 12°C, 23°C for transfer at 4°C, 12°C for transfer at 4°C, and a soil moisture content of 80% for 14 days. B, Statistical analysis of the survival rate of the samples in (A). C, Relative expression levels of *CCA1* in the WT after 12°C/23°C treatment and after 12°C/23°C/4°C transfer. D, Statistical analysis of the MDA content of the seedlings in (A). The data are presented as the means ± SEMs (n = 3 biological replicates). Different letters indicate significant differences (*P* < 0.05, one-way ANOVA, Tukey’s HSD test).

We also explore the influence of pre-treatment with moderate cold treatment (12°C) under dry and sharp cooling conditions, 3-week-old plants were exposed to either direct or pretreated moderate cold treatments along with simultaneous drought stress (Supplemental Figure S8, A and F). The findings demonstrated that moderate cold treatment significantly enhanced drought tolerance; survival rates were notably higher when combined with drought stress compared to drought stress alone (Supplemental Figure S8B). Additionally, a 2-day period of moderate cold pretreatment also improved plants ability to withstand both low temperatures and drought conditions; survival rates were significantly higher than those observed in plants treated solely with drought stress or direct dry cooling without any prior preparation (Supplemental Figure S8, A and B). The results of MDA content determination also verified the conclusion (Supplemental Figure S8D). The changes in *CCA1* expression were also investigated under different treatment conditions, and the results demonstrated that both medium-low temperature treatment and medium-low temperature pretreatment significantly enhanced the expression of the *CCA1* gene (Supplemental Figure S8E).

### The regulation of *P5CS1* and *OST1* expression by CCA1 is conserved across different species

Amino acid sequences of Arabidopsis CCA1 (AT2G46830) was used to perform BLAST searches in Phytozome v11.0 to identify putative *CCA1* genes throughout the plant kingdom. As a result, a total of 83 CCA1 genes were identifed from 64 plant species, including 39 dicots, 15 monocots, 5 algae, 3 bryophytes and 1 spikemoss (Selaginella moellendorfi). A rooted phylogenetic tree of CCA1 family was constructed based on multiple protein sequences alignment of above-mentioned 83 genes using Bayesian methods (Figure 8A). The constructed Bayesian tree was associated with overall high bootstrap values represented by colour gradient. Two major clades were found and they represent algae (in orange) and land plants. Within land plants, a large clade representing angiosperms was formed including two subclades specifc to monocot (in green) and dicot (in blue), respectively, together with the early-diverging angiosperm, Amborella (Figure 7A).

**Figure 8.**
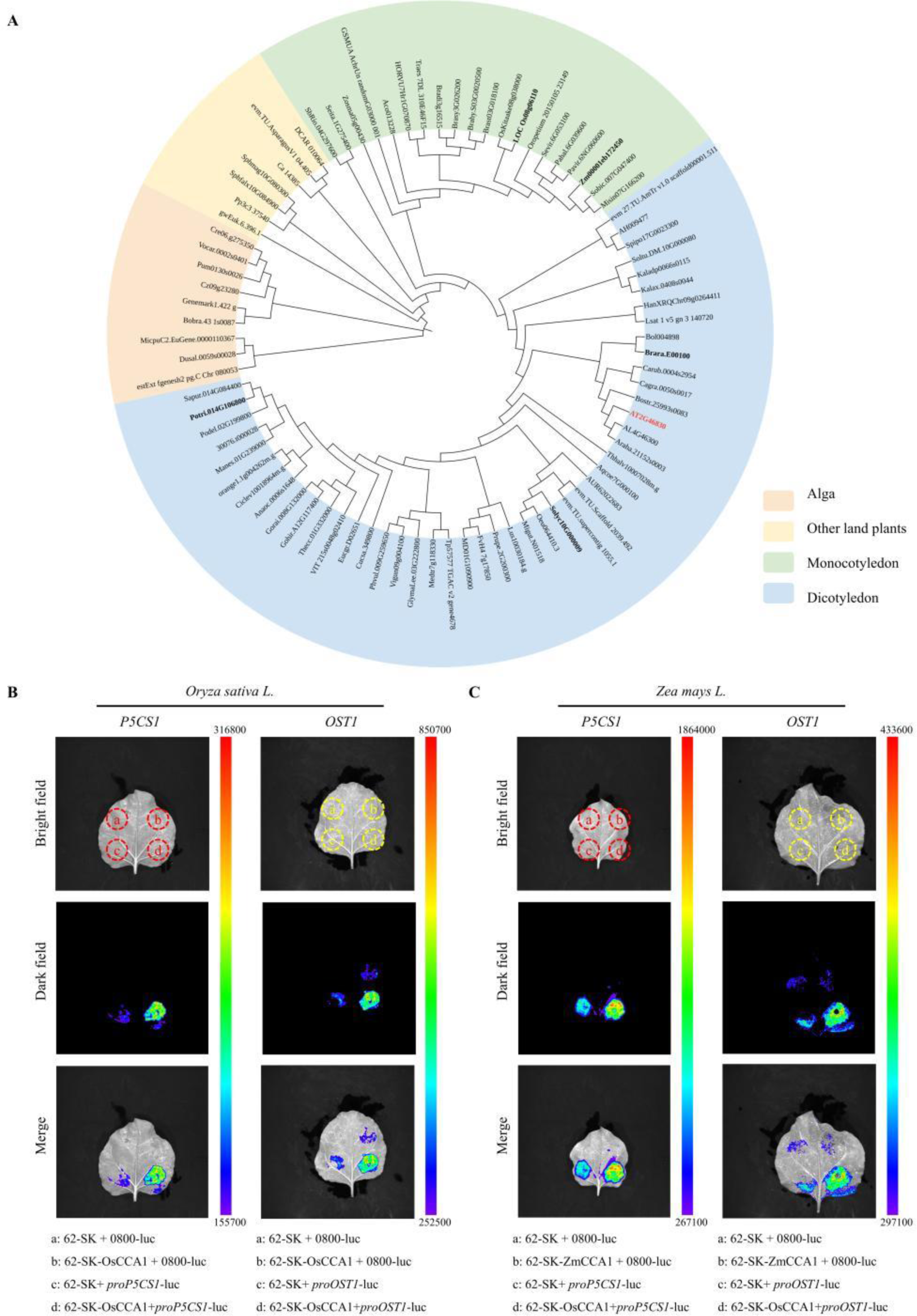
Evolutionary analysis of CCA1 regulation of drought tolerance in response to cold stress. A, Phylogenetic analysis of the *CCA1* genes in plants. A phylogenetic tree was constructed using the Bayesian method based on the amino acid sequences of 83 CCA1 members from 64 plant species. B, Luciferase reporter experiment of both OsCCA1 and the promoters of *OsP5CS1* and *OsOST1*. C, Luciferase reporter experiment of both ZmCCA1 and the promoters of *ZmP5CS1* and *ZmOST1*.

**Figure 9.**
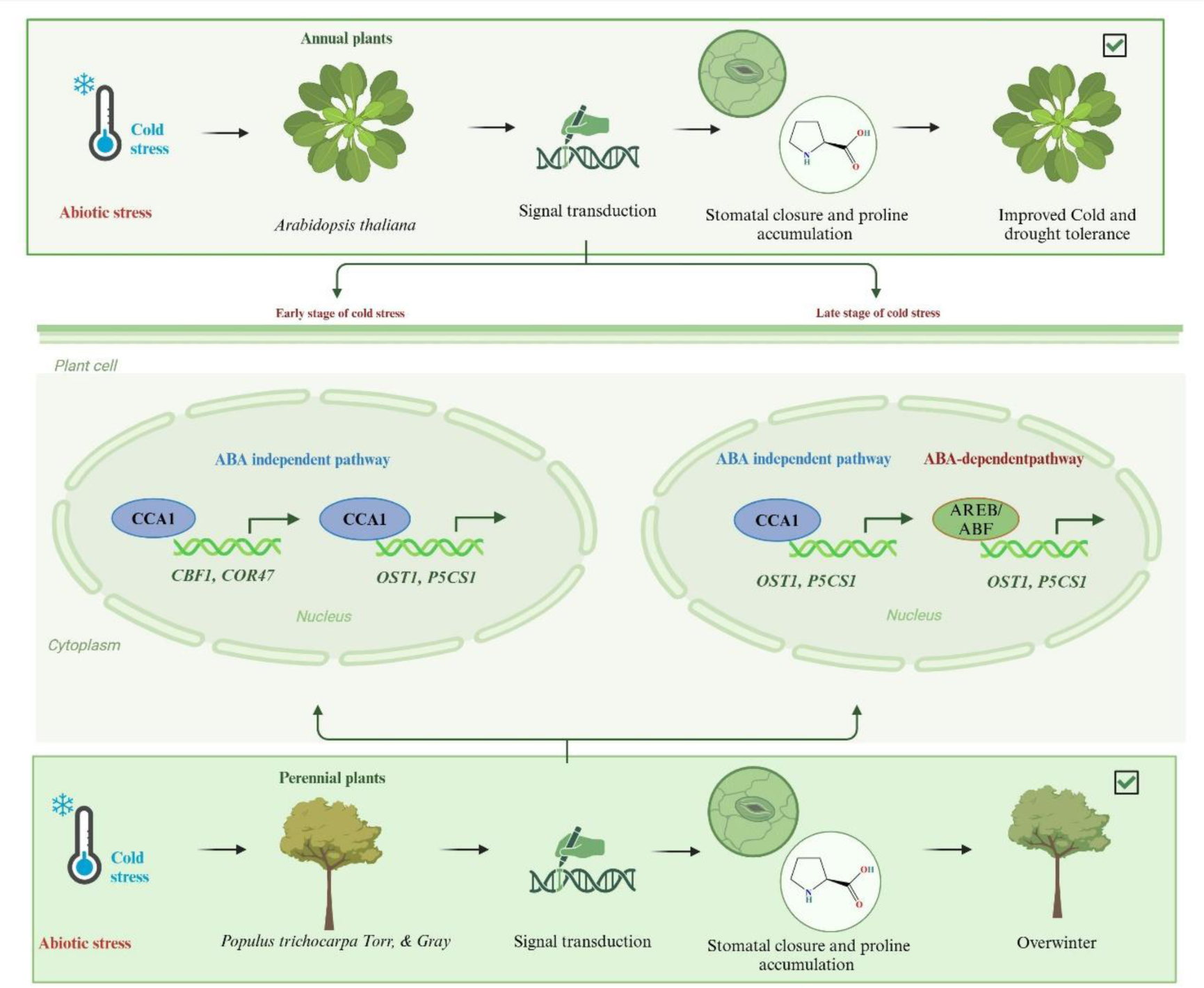
Moderate cold stress mediates high expression of *CCA1* through ABA-independent pathways and enhances drought resistance in *Arabidopsis* by targeting transient EE-acting elements in the *OST1* and *P5CS1* promoter regions.

We conducted a prediction of CCA1 binding elements within the promoter regions of *P5CS1* and *OST1* in different species, revealing a notable disparity in the abundance of these elements between their respective promoters in algae and land plants, more regulatory elements are abundant in terrestrial plants (Supplemental Figure S9, A and B). This suggests that *CCA1* may co-evolve with *OST1* and *P5CS1* in terrestrial plants, and participate in CCA1-mediated environmental adaptation. We also validated the transcriptional activation of *P5CS1* and *OST1* by CCA1 in rice (*Oryza sariva L.*) (Figure 7B), maize (*Zea miays L.*) (Figure 7C), and cabbage (*Brassica rapa var. glabra* Regel) (Supplemental Figure S10A), poplar (*Populus trichocarpa Torr, & Gray*) (Supplemental Figure S10B), tomato (*Solanum lycopersicum L.*) (Supplemental Figure S10C). The findings demonstrated that similar regulatory relationships were present in both monocotyledonous and dicotyledonous Arabidopsis plants; specifically, CCA1 was capable of inducing the transcription of *P5CS1* and *OST1*. These results imply that the mechanism through which CCA1 detects temperature fluctuations in terrestrial plants and enhances drought and cold resistance is conserved.

## Discussion

Temperature is a key environmental factor in plant physiology and ecology, which directly affects plant biochemical reactions, metabolic processes, as well as growth and development. Cold and drought stress are the two most common abiotic stresses that impede plant growth and development. These two stresses can occur simultaneously, especially in northwestern and northeastern China, posing significant constraints on agricultural productivity (Jiang et al. 2017; Wang et al. 2016). Multiple histological and genome-wide association analyses have demonstrated the convergence of transcripts, proteins, and metabolites in response to cold and drought stresses (Sharma et al. 2018). Moreover, both cold and drought stress induce cellular dehydration, thereby leading to overlapping signaling pathways such as stomatal closure and osmotic substance accumulation that contribute to cellular dehydration prevention (Kissoudis et al. 2014).

Current research on low temperature and drought stress has shown that pre-treatment of plants with drought stress can be effective in increasing tolerance to various abiotic stresses, including low temperature (Li et al. 2015; Hoffman et al. 2012; Saini et al. 2019; Guo et al. 2021). And also shown that plants exposed to non-lethal low temperatures for several days develop enhanced resistance to subsequent freezing stress (Guy 1990), which is related to the fact that non-lethal low temperatures induce the accumulation of large quantities of osmoregulatory substances that are also generally beneficial for drought resistance. Interestingly, our results showed that the growth status of Arabidopsis thaliana in the dry with moderate low temperature treatment was better than that of drought treatment alone but worse than that of moderate low temperature treatment alone, and the damage received by the combined treatment was also less than that of drought treatment alone but higher than that of moderate low temperature treatment alone (Figure 1). The results suggest that subjecting Arabidopsis to a cold treatment condition of 12 °C positively enhanced its drought tolerance. However, it was observed that the plants cold resistance did not improve under drought treatment, this is inconsistent with the results of previous studies (Saini et al. 2019). The reason behind this phenomenon may be attributed to our simultaneous treatment rather than pre-treatment. Our subsequent results also showed that plants first responded to cold stress while drought stress lagged behind when treated simultaneously (Figure 2). The previous findings have also demonstrated that pre-exposure to drought can enhance plants cold resistance by promoting the accumulation of ABA and other stress resistant substances (Li et al. 2015; Hoffman et al. 2012; Saini et al. 2019; Guo et al. 2021). This augmentation in cold tolerance is essentially attributed to the prior induction of drought stress in plants.The combination of our findings with the previous results leads us to believe that, in the context of compound stress, early response-induced stress is advantageous over delayed response-induced stress.

Based on the above understanding, we conducted a screening for transcription factors that exhibited rapid induction in response to cold stress specifically. The results demonstrated that *CCA1* exhibited specific induction in response to cold stress, while remaining unresponsive to drought stress. Furthermore, this induction was observed at an early stage during the cold stress treatment (Figure 2B; Supplemental Figure S2A). CCA1 is a transcription factor expressed in the morning and is closely related to the establishment of the circadian clock (Wang and Tobin 1998). And *CCA1* has also been reported to be involved in various abiotic stress responses. CCA1 can regulate the expression of *CBF1*, *CBF2* and *CBF3* in response to cold stress (Dong, Farré, and Thomashow 2011), and plants can coordinate their physiological and developmental responses to the environment through the regulation and feedback regulation of *CCA1* and *ABF3* (Liang et al. 2024). FBH1 can mediate the high-temperature signal to the circadian clock by directly targeting *CCA1* (Nagel, Pruneda-Paz, and Kay 2014). In this study, we have demonstrated that *CCA1* exhibits rapid cold stress perception and subsequently modulates stomatal closure by regulating *OST1* expression, as well as influences proline accumulation through the regulation of *P5CS1* expression, which significantly enhance plant drought resistance (Figure 3 and 4). Our results demonstrate that *cca1* mutant exhibit a loss in their capacity for cold preconditioning to enhance drought stress, while over-expression amplifies this ability, cold stress-induced stomatal closure and proline accumulation were masked in cca1 mutants (Figure 3). Over-expression of *OST1* and *P5CS1* can rescue the phenotype of increased stomatal opening and reduced proline accumulation of the *cca1* severally (Figure 5, B and H), and thus greatly compensate the cold-induced drought resistance of the mutant (Figure 5, A and F). Our results also suggested that CCA1 could directly bind to the promoters of *OST1* and *P5CS1* and positively regulates their transcription (Figure 4). The findings suggest that CCA1 plays a pivotal role as a transcription factor in mediating the effects of cold stress-induced drought stress resistance, while *OST1* and *P5CS1* act as downstream genes regulated by CCA1.

ABA is an important stress hormone involved in responses to many abiotic stresses, including cold and drought (Zhang, Zhu, et al. 2022; Shi, Ding, and Yang 2018a). ABA also regulates the expression of *OST1* and *P5CS1* (Zhang, Zhu, et al. 2022). To clarify whether drought stress resistance induced by cold stress is related to ABA accumulation. We monitored the association between ABA accumulation and stomatal opening under different treatment conditions using fluorescent staining techniques. The results showed that stomatal closure was earlier than ABA accumulation under cold treatment (Supplemental Figure S6). In ABA-deficient mutants, cold stress also regulates expression of *OST1* and *P5CS1* by inducing *CCA1* expression. These results suggest that cold stress enhances drought resistance through *CCA1* independently of ABA (Figure 6). Our results also show that this regulatory mechanism is conservative, and similar regulatory patterns exist in terrestrial dicotyledonous and monocotyledonous plants (Figure 8). In terrestrial plants, CCA1 is relatively conserved, and the promotors of *OST1* and *P5CS1* regulated by it also contain multiple CCA1 regulatory sites. These results suggest a possible co-evolution between them.

The simultaneous occurrence of cold and drought stress poses a challenge to plants in the middle and high latitudes, especially in the northwest and northeast regions of China, and has become an important factor limiting plant growth. In these areas, the gradual cooling in winter is usually accompanied by a decrease in rainfall and a gradual drying of the soil. Our results showed that moderate cold stress pretreatment could significantly improve plant tolerance to cold stress and drought stress. This may partly explain why overwintering plants are intolerant to extreme cooling or prolonged drought stress, but can tolerate seasonal extreme low temperatures and prolonged drought after gradual cooling in winter. This will provide a new target and theoretical support for the subsequent improvement of the wintering ability of crops and trees.

## Materials and Methods

### Transgene constructs

The full-length coding sequence (CDS) of *AtCCA1* was amplified from cDNA of *Arabidopsis thaliana* (Columbia-0) and cloned into the vector pENTRTM/SD/D-TOPO® (Invitrogen) and then transferred to the gateway-compatible pMDC32 binary vector and pMDC43-green fluorescent protein (GFP) vector using LR recombination (Life Technologies) (Smedley and Harwood 2015; Amack and Antunes 2020) to generate a vector for 35S-CCA1 and CCA1-GFP. These constructs were then transformed into Columbia-0 by Agrobacterium-mediated transformation using the floral-dip method (Purwantoro et al. 2023) to obtain *35S::CCA1* and *35S::CCA1-GFP* transgenic lines.

35S-CCA1 constructs were then transformed into *p5cs1*, *ost1* and *aba1* mutant by Agrobacterium-mediated transformation to obtain *35S::CCA1/ p5cs1*, *35S::CCA1/ ost1* and *35S::CCA1/ aba1* transgenic lines. All primers used to generate transformation constructs are listed in table S1.

### Plant materials and growth conditions

The *Arabidopsis thaliana* Columbia-0 ( ecotype Col-0) was used as the wild-type (WT). The T-DNA insertion mutants *cca1* (SALK_146072C), *p5cs1* (SALK_063517C), *ost1* (SALK008068C) and aba1 (SALK_027326) were obtained from the Arabidopsis Biological Resource Center (ABRC). Plants were routinely grown in soil mixture containing a 1:1 ratio of nutrient soil and vermiculite. The plants were maintained at 23°C and 60% humidity under fluorescent light (152 mmol photons·m^-2^·s^-1^) with a 16 h light/8 h dark cycle. Prior to in vitro culture, seeds of Arabidopsis were surface sterilized with 75% ethanol for 1 min, 3 times in total, and then washed with 95 alcohol for 1 minute, also 3 times. The seeds were then washed 4-6 times with sterile distilled water and sown on MS agar plates (Sigma, St. Louis, MO) supplemented with 0.8% sucrose. Following stratification at 4°C for 2 days in the dark, the plates were sealed and incubated at 23°C in a chamber under fluorescent light (152 mmol photons·m^-2^·s^-1^) with a 16 h light/8 h dark cycle. For tobacco (*Nicotiana benthamiana*) plants, seeds were sown in a steam-sterilized compost soil mix (nutrient soil and vermiculite, 1:1, v/v/v). The tobacco plants were raised in a growth chamber at 25 ± 1°C under the conditions described above.

### Stress treatment

For control group, 3-week-old Arabidopsis seedlings were exposed to 23℃ and soil moisture content 80% for 14 days. For cold group, 3-week-old Arabidopsis seedlings were exposed to 12℃ and soil moisture content 80% for 14 days. For drought group, 3-week-old Arabidopsis seedlings were exposed to 23℃ and then drought treatment was started until soil moisture content 40% for 14 days. For Cold and Drought group, 3-week-old Arabidopsis seedlings were exposed to 12℃ for 1day and then drought treatment was started until soil moisture content 40% for 14 days.

### Relative water content measurements

For measuring the relative water content (RWC) (González and González-Vilar 2001), aerial parts were collected from 3-week-old Arabidopsis seedlings of each plant lines (n = 30) at 30, 60, 90, 120 and 150 min after exposure different treatment groups, and its fresh weights were measured. Leaves were hydrated for 12 h in distilled water to identify their turgid weight (TW) and then dried at 105 °C for 5 min and at 80 °C until constant weight for dry weight (DW). RWC was calculated as follows: RWC (%) = (FW − DW)/(TW − DW) × 100%.

### Proline content measurements

For measuring the proline content (Senthilkumar, Amaresan, and Sankaranarayanan 2021), aerial parts were collected from 3-week-old Arabidopsis seedlings of each plant lines after exposure different treatment groups for 14 day. One hundred milligrams of frozen plant material was homogenized in 1.5 ml of 3% sulfosalicylic acid and centrifuged. One hundred microliters of the extract was reacted with 2 ml glacial acetic acid and 2 ml acid ninhydrin (1.25 g ninhydrin warmed in 30 ml glacial acetic acid and 20 ml 6 M phosphoric acid until dissolved) for 1 h at 100 °C, and the reaction was then terminated in an ice bath. The reaction mixture was extracted with 1 ml toluene and measured at 520 nm. The proline content was calculated according to a standard curve.

### Malondialdehyde content measurements

For measuring the malondialdehyde (MDA) content (Tulkova and Kabashnikova 2022; Morales and Munné-Bosch 2019), aerial parts were collected from 3-week-old Arabidopsis seedlings of each plant lines after exposure different treatment groups for 14 day. Leaves were homogenized in 5% (w/v) trichloroacetic acid (TCA) and centrifuged at 3000× g for 10 min. Supernatants were collected and reacted with an equal volume of 0.67% (w/v) thiobarbituric acid (TBA) in a boiling water bath for 30 min. After cooling, the mixture was centrifuged at 3000× g for 10 min. The absorbance of supernatant was measured at 532 nm and corrected for nonspecific turbidity by subtracting the absorbance at 600 nm and 450 nm.

### Relative electrical conductivity measurements

For measuring the relative electrical conductivity (Cha et al. 2021), aerial parts were collected from 3-week-old Arabidopsis seedlings of each plant lines after exposure different treatment groups for 14 day. Leaf samples were detached from each plant line and placed into tubes containing 20 mL of deionized water (S0). Following shaking for 12 h at room temperature, the conductivity of the samples was measured (S1). After autoclaving for 15 min, the tubes were shaken for 1 h and the conductivity was measured (S2). Electrolyte leakage of each sample was calculated using the formula S1 – S0/S2 – S0.

### RNA extraction and qRT-PCR

Total RNA extraction from plants was isolated with the RNAprep Pure Plant Kit (TIANGEN DP432). cDNA was synthesized using 1 µg of total RNA by utilizing the FastQuant RT Kit (with gDNase) (TIANGEN KR106). Synthesized cDNAs were then quantified by qRT‒PCR with PerfectStart® Green qPCR SuperMix (TRAN AQ601) using the CFX96 Real-Time PCR System (Bio-Rad). Actin 8 (AT1G49240) was used as a reference gene in Arabidopsis. The PCR conditions used were 94 °C for 30 s followed by 40 cycles at 94 °C for 5 s and 60 °C for 30 s. The results were repeated three times. Each sample and statistical analysis were performed using a standard curve method. The oligomers for qPCR are shown in Table S1.

### Subcellular localization assays

The 35S-GFP (control vector) and 35S-*AtCCA1*-GFP plasmids were transfected into were subsequently transformed into Agrobacterium tumefaciens GV3101 cells, which were subsequently co-infiltrated into tobacco (*Nicotiana benthamiana*) leaves. After 48-72 hours, the GFP fluorescence signal (excitation and emission wavelengths set to 488 nm) was detected using a confocal microscope (LSM880 META, Zeiss, Germany) to determine the protein.

### Dual luciferase reporter assay

The approximately 2-kb promoter fragment of *AtP5CS1*, *AtOST1, OsP5CS1*, *osOST1*, *ZmP5CS1*, *ZmOST1*, *BrP5CS1*, *BrOST1*, *PtP5CS1*, *PtOST1*, *SyP5CS1*, and *SyOST1*, were synthesized and cloned into the pGREEN II 0800-LUC vector digested by KpnI and BamHI, generating the reporter vectors. The full-length CDS of *AtCCA1*, *OsCCA1*, *ZmCCA1*, *BrCCA1*, *PtCCA1*, *SyCCA1* was amplified and inserted into vector pGREEN II 62-SK digested by BamHI and KpnI to generate effector vector. The empty pGREEN II 62-SK and pGREEN II 0800-LUC vector was used as negative control. Equal amounts of effector and reporter vectors were co-transformed into *Nicotiana benthamiana* and incubated in the 12light/12dark at 28 °C for 3d. The *N. benthamiana* leaves were sampled, and signals were visualized using a NightSHADE LB 985 in vivo Plant Imaging System (BERTHOLD). IndiGO™ software was used to digitize the experimental results. The luciferase (LUC) activity was measured with the Dual-Luciferase Reporter Assay System Kit (TransDetect® FR201-01-V2) using the TriStar2 Multimode Reader LB942 (Berthold Technologies), with the Renilla luciferase (REN) gene driven by the CaMV 35S promoter as an internal control. Relevant primer sequences are listed in table S1.

### Electrophoretic mobility shift assay (EMSA)

The coding region of *CCA1* was inserted into the pMAL-c2x vector at the BamHI and SalI sites. The plasmid was expressed in Escherichia coli Rosetta 2(DE3) Chemically Competent Cells (Wei di, Shanghai, China) for expression of recombinant MBP-AtCCA1 protein, followed by purification using Anti-MBP Magnetic Beads (Beyotime, Shanghai, China). DNA probes of 40-50 bp length were chemically synthesized and labeled with biotin (Sangon Biotech, China). The binding reactions and DNA gel-shift assays were performed with the LightShift Chemiluminescent EMSA Kit (Thermo Fisher Scientific, 20148). Membranes were imaged using a Tanon 4800 multi Chemiluminescence Imager device (Tanon). Relevant probe sequences are listed in table S1.

### Chromatin immunoprecipitation assays (ChIP assays)

The ChIP-qPCR assays were conducted using the ChIP kit (EpiQuikTM Plant ChIP Kit, Sigma) and the corresponding protocol. 7 to 10-day-old seedlings Arabidopsis of WT and *35S::CCA1-GFP* transgenic lines to perform ChIP-seq. The samples were cross-linked with 1% formaldehyde. Glycine (2 M) was added to the cross-linking buffer to stop the reaction. Plant samples were then removed from the buffer and frozen in liquid nitrogen to homogenize and release the nucleus. The remaining steps, such as nucleus isolation, protein immunoprecipitation, DNA elusion, and DNA purification, were performed according to the protocols provided with the kits. The purified DNA and the DNA before immunoprecipitation (input) were used to conduct qPCR tests. For qPCR tests, PerfectStart® Green qPCR SuperMix (TRAN AQ601) was used for qPCR using the CFX96 Real-Time PCR System (Bio-Rad). The primers used in the qPCR test are summarized in Table 1. The final data were analysed by the following step (Mo et al. 2022). The results were normalized to input DNA using the equation 2^(Ct input − Ct ChIP) × 0.1.

### Microscale thermophoresis assay

A NanoTemper Monolith biomolecular interaction detector was used to detect protein-nucleic acid interactions and binding activities. When detecting protein-nucleic acid interactions, CCA1, as a target, connects to the GFP fluorescent tag, and, the nucleotide sequence as ligands. Each assay consisted of 16 reaction mixtures with the ligand concentration diluted in half in turn. NT.115 ConFigure the channel as Nano BLUE/RED, and detection was performed when the MST power was set to 40%. In the inhibition binding experiment, the target and another binding protein were incubated in the dark at room temperature for several minutes, and then, a gradient dilution of the ligand was added for reaction detection. Finally, MO. Affinity Analysis v2.3 was used for the data analysis.

### Stomatal Bioassay

For the determination method of stomatal parameters, refer to (Chen et al. 2010; Chen et al. 2016). The specific operation steps are as follows: cut the smooth 4-week Arabidopsis rosette leaves and soak them in a transparent glass Petri dish equipped with a closure buffer (20 mM KCl, 5 mM MES, 1 mM CaCl_2_ adjusted to pH 6.1 with K(OH) 2, 0.003% Silwet-77). The control group was cultured in a 21 °C light incubator for 2 h, and the experimental group was treated in a 4 °C light incubator for 2 h with a light intensity of 100 μ Mol m^-2^ S^-1^ PAR to fully open the stomatal pores. The mesophyll cells were removed, and pictures of guard cells were taken using a microscope. The images were collected by an Axio Observer inverted microscope (Zeiss, Oberkochen, Germany) and observed for 120 minutes. The whole determination process of stomatal pore parameters was 100 μMol m^-2^ S^-1^ par, which was used to avoid dark-induced stomatal closure and affect the experimental results. During the observation, image acquisition was carried out every 5 minutes, and the stomatal opening and other parameters were measured by ImageJ software. Thirty to forty stomata were measured at each time point (technical repetition). The final results were calculated by 3 biological repetitions.

### Immunofluorescence and confocal microscopy

Immunofluorescence analysis of stomatal closure in Arabidopsis leaves was performed as previously described by Andème-Onzighi et al. (Andème-Onzighi et al. 2002) with a few modifications. Whole 3-week-old Arabidopsis leaves were vacuum-infiltrated for 2 h and then fixed overnight at 4°C in 50 mM sodium PBS, pH 7.2, containing 4% (v/v) paraformaldehyde (EMS), 0.2% (v/v) glutaraldehyde (Sigma-Aldrich), and 2% (v/v) EDC (Thermo Scientific), an essential step allowing the fixation of ABA in situ, via crosslinking by its carboxyl group (Peng et al. 2006; Sotta et al. 1985; Sossountzov et al. 1986). Fixed stomatals were washed three times with PBS and then were simultaneously blocked and permeabilized with 3% (w/v) nonfat dried milk solution in 0.01 M PBS (PBS-milk), pH 7.2, containing 0.1% cellulose, 0.1% pectinase, and 0.1% Triton for 1 h. Permeabilized root tips were washed three times with PBS and incubated with 0.25 µg anti-ABA polyclonal antibodies (rabbit anti-ABA, C1, AS09446; Agrisera) overnight at 4°C (Turecková, Novák, and Strnad 2009; Hradecká et al. 2007). The stained leaves were washed and transferred to a 400-fold dilution of secondary antibody goat anti-rabbit IgG-conjugated to Alexa Fluor 488 (excitation, 488 nm; emission, 505 to 530 nm) (Invitrogen, Molecular Probes) for 2 h at room temperature. The stained leaves were washed with PBS and mounted with Citifluor AF1 (Ted Pella) and then imaged by confocal microscopy (LSM880 META, Zeiss, Germany) to determine the proteins.

### Structural prediction

Single protein structures are derived from the AlphaFold Protein Structure Database. We use AlphaFold 3 (AlphaFold Server) to predict protein-DNA interactions. The proteins use all amino acid residues, the *P5CS1* promoter sequence are “ 5′-attgaaaaatatctaaatactgt-3′ “ and “5′-acagtatttagatatttttcaat-3′ ”, and the *OST1* promoter sequence are “ 5′-gctagccaacaatatctcgacat -3′ “ and “5′-atgtcgagatattgttgttggctagc-3′ ”. AlphaFold 3 provides a graph of expected position error, where the darker the color, the smaller the error and the higher the confidence.

### Statistical analysis

Statistical analyses were performed using Excel and SPSS software. Analysis of variance was used to compare significant differences based on Students t test at significance levels of *P* < 0.05.

## AUTHOR CONTRIBUTION

M-XC and Y-GL designed the experiments. XY, YL, Z-CJ, ML, X-XH, S-QH and X-LS performed the experiments and analysis. M-XC, F-YZ, BG, D-YZ, and Y-GL drafted the article. Y-GL and M-XC critically reviewed and revised the manuscript.

## ACKNOWLEDGEMENTS

This work was supported by the National Natural Science Foundation of China (NSFC32272132, U2106230 and 32001452) and Shandong Qingchuang team Fund.

## COMPETING INTEREST

There are no competing interests to declare.

